# Identification of candidate causal *cis*-regulatory variants underlying electrocardiographic QT interval GWAS loci

**DOI:** 10.1101/2024.03.13.584880

**Authors:** Supraja Kadagandla, Ashish Kapoor

**Author notes:** Corresponding author: Ashish Kapoor, Ph.D., Center for Human Genetics, Institute of Molecular Medicine, McGovern Medical School, University of Texas Health Science Center at Houston, 1825 Pressler Street, Room SRB 530C, Houston, TX 77030, USA, (T), (E).

## Abstract

Identifying causal variants among tens or hundreds of associated variants at each locus mapped by genome-wide association studies (GWAS) of complex traits is a challenge. As vast majority of GWAS variants are noncoding, sequence variation at *cis*-regulatory elements affecting transcriptional expression of specific genes is a widely accepted molecular hypothesis. Following this *cis*-regulatory hypothesis and combining it with the observation that nucleosome-free open chromatin is a universal hallmark of all types of *cis*-regulatory elements, we aimed to identify candidate causal regulatory variants underlying electrocardiographic QT interval GWAS loci. At a dozen loci, selected for higher effect sizes and a better understanding of the likely causal gene, we identified and included all common variants in high linkage disequilibrium with the GWAS variants as candidate variants. Using ENCODE DNase-seq and ATAC-seq from multiple human adult cardiac left ventricle tissue samples, we generated genome-wide maps of open chromatin regions marking putative regulatory elements. QT interval associated candidate variants were filtered for overlap with cardiac left ventricle open chromatin regions to identify candidate causal *cis*-regulatory variants, which were further assessed for colocalizing with a known cardiac GTEx expression quantitative trait locus variant as additional evidence for their causal role. Together, these efforts have generated a comprehensive set of candidate causal variants that are expected to be enriched for *cis*-regulatory potential and thereby, explaining the observed genetic associations.

## Introduction

Despite the challenges in elucidating the molecular mechanisms underlying genome-wide association studies (GWAS) loci (1, 2), as an unbiased screen, GWAS have made immense contribution towards understanding the genetic architecture of complex traits (3). However, GWAS maps and identifies positional markers or variants, not causal variants nor genes. Unlike Mendelian disorders, where causal variant detection and gene identification are synonymous, functional follow up of GWAS mapping of complex traits requires substantial work to uncover the genes and variants involved (4). Nevertheless, a widely accepted molecular hypothesis of complex trait variation is sequence variation largely, but not exclusively, at *cis*-regulatory elements affecting transcriptional variation of specific genes (5-8). Mapping of complex traits by GWAS has paralleled annotation of putative functional elements regulating gene expression (9) in the human genome by efforts such as the ENCODE (10) and NIH RoadMap Epigenomics (11) projects, and expression quantitative trait locus (eQTL) mapping (12, 13) of variants associated with population-level gene expression variability by efforts such as the GTEx (14) project. These large-scale efforts have provided at least the initial set of tools to test the *cis*-regulatory mechanistic hypothesis for GWAS signals in the noncoding genome using a variety of empirical wet-lab, statistical and bioinformatic approaches (2).

Among various functional annotations of the human genome (10, 11) that can predict *cis*-regulatory elements (9), chromatin accessibility, determined by DNase I sequencing (DNase-seq) (15) or Assay for Transposase Accessible Chromatin using sequencing (ATAC-seq) (16), and visualized as DNase peaks and ATAC peaks, is by far the best predictors of *cis*-regulatory elements and universal hallmark of all types of *cis*-regulatory elements (17). Furthermore, numerous common disease-associated variants can be identified in putative *cis*-regulatory elements marked as open chromatin genome-wide (6), and such element identification and GWAS variants enrichment is improved when the cell-type relevant to a trait or disease is used (18). Following these observations and the widely accepted *cis*-regulatory mechanistic hypothesis underlying GWAS signals, our goal here was to use bioinformatic approaches to integrate electrocardiographic QT interval GWAS variants (19) with DNase-seq and ATAC-seq based genome-wide maps of cardiac open chromatin regions to identify candidate causal *cis*-regulatory variants, and further prioritize them for causality if they are known to act as eQTLs in cardiac tissues.

Here, using bioinformatic approaches we report identification of candidate causal *cis*-regulatory variants at 12 selected GWAS loci for electrocardiographic QT interval. Among all known QT interval GWAS loci, we limit to a dozen loci with relatively higher effect sizes and encompassing genes known to directly or indirectly influence cardiomyocyte action potential duration or excitation-coupling. At each locus, we use linkage disequilibrium (LD) to expand the set of associated candidate variants. In parallel, using ENCODE DNase-seq and ATAC-seq datasets from multiple adult cardiac left ventricle tissue samples, we generate genome-wide maps of open chromatin regions. Filtering associated candidate variants for overlap with cardiac left ventricle open chromatin regions, we identify candidate causal *cis*-regulatory variants, and further prioritize them if they colocalize (i.e., same variant) with GTEx cardiac eQTLs.

## Methods

### QT interval GWAS loci

Among several dozens of associated loci identified by QT interval GWAS (20), we selected 12 loci (*ATP1B1, CNOT1, KCNH2, KCNJ2, KCNQ1, LIG3, LITAF, NOS1AP, PLN, RNF207, SCN5A* and *SLC81A1*; named based on the gene closest to the sentinel variant) with relatively larger effect sizes that were reported for the first time in 2009 (21, 22), and have been replicated since then in multiple studies (19, 20).

### Definition of the target GWAS locus

At each locus, using the GWAS summary statistics (19), the most proximal and the most distal variants meeting the genome-wide significance threshold (*P*<5×1o-8) and within ±500kb of the top GWAS hit were identified. The target locus was then defined using HapMap Phase II variants-based recombination hotspot (23) immediately upstream and downstream of the most proximal and most distal genome-wide significant variant (GWS), respectively.

### Identification of high LD variants

Data Slicer tool was used to get subsections of 1000 Genomes Phase 3 variant call format (VCF) files (24) based on the genomic coordinates of each target GWAS locus. At each locus, biallelic variants observed in the five EUR ancestry subpopulations (CEU, FIN, GBR, IBS and TSI) with minor allele frequency (MAF) above 1% and in high LD (*r*^2^>0.9) with any of the GWS variants were identified using VCFtools (25).

### Identification of open chromatin regions

Filtered binary alignment map (BAM) files from paired-end DNase-seq (15) and ATAC-seq (16) in human adult heart left ventricle tissue samples were downloaded from the ENCODE project (10) portal. Integrity of the BAM files was checked using SAMtools (26) stats function and matched with the same available at the ENCODE project portal. Duplicate reads were flagged and removed from DNase-seq BAM files and reads with low mapping quality (MAPQ<20) were removed from ATAC-seq BAM files using SAMtools. Next, these processed BAM files were converted to BEDPE (browser extensible data (BED) paired-end) using BEDtools bamToBed utility, where only one alignment from paired-end reads is reported. The two cut positions for each alignment were extracted from the BEDPE files separately to convert them into BED files. To account for the 9-bp duplication of the target site by Tn5 transposase in ATAC-seq library generation (16, 27), + strand reads were offset by +4 bp and - strand reads were offset by -5 bp to identify the two cut positions for each alignment in ATAC-seq. Next, ATAC-seq and DNase-seq peaks were identified in each sample using MACS2 (28) with the option “--nomodel --shift -50 --extsize 100 --keep-dup all”, and sample-level open chromatin regions were identified as 600-bp regions centered at the summits of MACS2 called peaks. To combine and merge overlapping open chromatin regions across samples, BEDtools (29) multiinter function was used to filter out complete or partial nonoverlapping regions that were observed only in any one sample, and the remaining regions were combined using BEDtools merge function.

### Identification of variants overlapping open chromatin regions

Trait associated variants overlapping human cardiac open chromatin regions were identified by generating BED files-based custom tracks in the UCSC Genome Browser (30) for the two, and using the Table Browser tool (31) to intersect the two custom tracks.

### Overlap with GTEx eQTLs

Single-tissue *cis*-eQTL (12, 13) data for left ventricle and atrial appendage were downloaded from GTEx (14) project portal (version 8). Trait associated variants overlapping open chromatin regions were assessed for being listed as an eQTL in the significant variant-gene pairs dataset.

### Genome build and conversions

GWAS summary statistics reported variants on the hg18 build. 1000 Genomes Phase 3 VCF files reported variants on the hg19 build. Both of these were converted to hg38 build using LiftOver tool in the UCSC Genome Browser. Filtered BAM files from the ENCODE project and GTEx single-tissue *cis*-eQTL data (version 8) were on the hg38 build. All genome coordinates presented here are on the hg38 build.

## Results

### Selection of QT interval GWAS loci

The most recent large-scale meta-analysis GWAS of QT interval, largely from European ancestry populations, published in 2022 (20) reported identification of dozens of associated loci, including replication of all major previously reported loci (19, 21, 22, 32). The first GWAS for QT interval was reported in 2006 and identified a single locus near *NOS1AP* (32). As samples size increased, two reports of QT interval GWAS in 2009 (21, 22) identified a dozen loci (*ATP1B1, CNOT1, KCNH2, KCNJ2, KCNQ1, LIG3, LITAF, NOS1AP, PLN, RNF207, SCN5A* and *SLC81A1*; named based on the gene closest to the sentinel variant), including the *NOS1AP* locus. Since then, these dozen loci have been replicated and have remained as the loci with relatively larger effects on variance explained in multiple studies (19, 20). Of all known QT interval GWAS loci, in the current study we focus on these dozen high impact loci. Another reason for limiting to these dozen loci is that these harbor genes encoding ion channels and their associated regulatory proteins that directly or indirectly influence cardiomyocyte action potential duration and/or excitation-contraction coupling, including *KCNQ1, KCNH2* and *SCN5A* that are known to underlie long QT, short QT and Brugada syndromes (33, 34).

### Identification of associated candidate variants based on high LD

We have built this study on a QT interval meta-analysis GWAS (19), performed largely in European ancestry populations, which used ∼2.5 million HapMap variants (23). To expand the set of associated variants, we identified all common variants observed in the 1000 Genomes project (24) EUR populations that are in high LD with the HapMap-based GWS variants. Starting with the GWAS summary statistics, we first identified the most proximal and most distal GWS variant within ±500 kb of the sentinel variant at each locus.

Recombination hotspot (23) immediately upstream and downstream of the most proximal and most distal GWS was used to define the boundaries of the associated locus (Table 1). Within this defined interval, we identified all 1000 Genomes Phase 3 variants observed in the five EUR populations that are common (MAF>1%; 503 subjects) and in high LD (*r*^2^>0.9) with any of the HapMap-based GWS variants. This led to a significant increase in the number of associated candidate variants at all loci (Table 1). Across the 12 loci, there were 1,401 HapMap-based GWS variants, which increased to 3,482 after including the 1000 Genomes Phase 3 common high LD variants (Table 1, Supplemental Datasets S1 to S12).

**Table 1:** QT interval associated candidate variants at each GWAS locus.

| Locus | Genome Position | #GWS Variants | #Common and High LD Variants | #Total Candidate Variants |
| --- | --- | --- | --- | --- |
| *ATP1B1* | chr1:168692142-169556075 | 102 | 222 | 324 |
| *CNOT1* | chr16:58329126-58835016 | 99 | 198 | 297 |
| *KCNH2* | chr7:150782497-151028636 | 94 | 139 | 233 |
| *KCNJ2* | chr17:70202991-70659990 | 84 | 116 | 200 |
| *KCNQ1* | chr11:2283323-2625391 | 30 | 68 | 98 |
| *LIG3* | chr17:34654986-35295756 | 38 | 37 | 75 |
| *LITAF* | chr16:11550985-11667258 | 26 | 108 | 134 |
| *NOS1AP* | chr1:161979398-162475347 | 354 | 422 | 776 |
| *PLN* | chr6:118011664-118730533 | 445 | 577 | 1022 |
| *RNF207* | chr1:6072008-6285075 | 14 | 43 | 57 |
| *SCN5A* | chr3:38310078-38822897 | 59 | 62 | 121 |
| *SLC8A1* | chr2:40107722-40559057 | 56 | 89 | 145 |

### Generation of cardiac left ventricle open chromatin maps

Open chromatin or increased chromatin accessibility is a universal hallmark of *cis*-regulation (17). Following the *cis*-regulatory mechanistic hypothesis for GWAS signals in the noncoding genome (5-8), an expectation is that majority of the causal variants will lie in phenotype-relevant open chromatin regions. In order to filter the associated candidate variants (see below), we first generated and combined open chromatin maps from human cardiac left ventricle tissue samples based on publicly available ENCODE (10) DNase-seq (15) and ATAC-seq (16) datasets, separately. There are eight and 15 human adult cardiac left ventricle tissues-based DNase-seq and ATAC-seq data, respectively, available from the ENCODE project (Supplemental Table S1). Of these one DNase-seq and nine ATAC-seq data are flagged for not meeting the ENCODE project’s standards at various levels, including low read coverage, library complexity and enrichment, and were excluded from further analyses (Supplemental Table S1). ENCODE project’s uniform pipeline for DNase-seq experiments at the post alignment filtering step removes reads with low mapping quality, but only marks duplicate reads. On the other hand, ENCODE project’s uniform pipeline for ATAC-seq experiments at the post alignment filtering step in multimapping mode removes duplicate reads, but not reads with low mapping quality. Therefore, filtered aligned BAM files for the remaining (seven DNase-seq and six ATAC-seq) were processed to remove PCR and/or optical duplicate reads (DNase-seq) or to remove low mapping quality reads (MAPQ<20; ATAC-seq) (Table 2).

**Table 2:** Processing of human adult left ventricle DNase-seq and ATAC-seq experiments for open chromatin peak calling.

| Assay | ENCODE Experiment ID | ENCODE BAM File ID | # Reads in ENCODE BAM | # Reads Post-processing | # MACS2-called Peaks | # Merged Open Chromatin Regions |
| --- | --- | --- | --- | --- | --- | --- |
| DNase-seq | ENCSR299QGI | ENCFF631HMZ | 193331564 | 142173006 | 291544 | 318334 |
| DNase-seq | ENCSR395HAE | ENCFF154ILG | 189236690 | 73211960 | 233919 |  |
| DNase-seq | ENCSR747SEU | ENCFF041ZOJ | 200256974 | 77074876 | 256470 |  |
| DNase-seq | ENCSR222CLC | ENCFF284XQW | 200569792 | 152422520 | 311689 |  |
| DNase-seq | ENCSR805EGZ | ENCFF221NLC | 125331996 | 115251966 | 199087 |  |
| DNase-seq | ENCSR598RVJ | ENCFF682TBE | 180874544 | 63724500 | 333492 |  |
| DNase-seq | ENCSR735NTV | ENCFF558NDU | 228946790 | 188384988 | 281851 |  |
| ATAC-seq | ENCSR286STX | ENCFF440WVI | 210452200 | 205323804 | 240542 | 188675 |
| ATAC-seq | ENCSR899YEP | ENCFF771XJN | 51252858 | 49792344 | 116368 |  |
| ATAC-seq | ENCSR310RJN | ENCFF804KIW | 139081990 | 135656876 | 198854 |  |
| ATAC-seq | ENCSR846VPV | ENCFF456PVX | 34141762 | 33316588 | 155876 |  |
| ATAC-seq | ENCSR925LGW | ENCFF827WPC | 249602824 | 243734600 | 259662 |  |
| ATAC-seq | ENCSR399OSE | ENCFF160LDI | 305801460 | 298336596 | 293473 |  |

Next, we performed peak calling with MACS2 (28) in the seven DNase-seq and six ATAC-seq datasets to identify varying number of peaks across the samples (199,087-333, 492 in DNase-seq; 116,368-293,473 in ATAC-seq). We defined sample-level open chromatin regions as 600-bp regions centered at the identified summits within peaks.

Complete or sub-open chromatin regions observed only in any one sample were excluded, and the remaining observed in any two or more samples, were combined to generate 318,334 and 188,675 merged open chromatin regions based on DNase-seq (Supplemental Dataset S13) and ATAC-seq (Supplemental Dataset S14), respectively, in human adult left ventricle tissue (Table 2).

### Filtering of associated candidate variants based on left ventricle open chromatin overlap

All associated candidate variants across the 12 GWAS loci were assessed for overlap with the left ventricle merged open chromatin regions derived from DNase-seq and/or ATAC-seq experiments. Across the 12 GWAS loci, of the 3,482 candidate variants (range: 57-1,022 per locus), 369 variants overlap DNase-seq based open chromatin regions and 317 variants overlap ATAC-seq open chromatin regions, for a total of 476 unique variants (range: 10-124 per locus) overlapping left ventricle open chromatin regions (Table 3; Supplemental Datasets S1 to S12); these are considered as candidate causal *cis*-regulatory variants underlying the GWAS signals. With the goal to further prioritize/rank them for causality and functional studies, we assessed if these variants are known to act as *cis*-eQTLs, thereby supporting their regulatory potential. Based on GTEx (14) single-tissue *cis*-eQTL (12, 13) data, of the 476 open chromatin overlapping associated candidate variants, 243 variants (eVariants) are known to be underlying an eQTL signal in human cardiac left ventricle and/or atrial appendage tissue (Table 3; Supplemental Datasets S1 to S12).

**Table 3:** Candidate variants overlapping cardiac open chromatin and acting as cardiac eQTLs.

| Locus | #Candidate Variants | #Variants Overlapping LV DNase-seq OCRs | #Variants Overlapping LV ATAC-seq OCRs | #Variants Overlapping LV OCRs | #Overlapping Variants as LV eVariants | #Overlapping Variants as AA eVariants | #Overlapping Variants as LV/AA eVariants |
| --- | --- | --- | --- | --- | --- | --- | --- |
| *ATP1B1* | 324 | 23 | 31 | 41 | 12 | 7 | 13 |
| *CNOT1* | 297 | 32 | 25 | 39 | 29 | 28 | 30 |
| *KCNH2* | 233 | 52 | 28 | 56 | 46 | 39 | 47 |
| *KCNJ2* | 200 | 7 | 7 | 10 | 0 | 0 | 0 |
| *KCNQ1* | 98 | 16 | 13 | 19 | 6 | 6 | 6 |
| *LIG3* | 75 | 9 | 6 | 10 | 9 | 9 | 9 |
| *LITAF* | 134 | 21 | 16 | 22 | 2 | 0 | 2 |
| *NOS1AP* | 776 | 72 | 69 | 97 | 56 | 26 | 56 |
| *PLN* | 1022 | 96 | 75 | 124 | 23 | 69 | 69 |
| *RNF207* | 57 | 13 | 16 | 18 | 4 | 6 | 6 |
| *SCN5A* | 121 | 14 | 14 | 18 | 4 | 1 | 5 |
| *SLC8A1* | 145 | 14 | 17 | 22 | 0 | 0 | 0 |

OCR: open chromatin region; LV: left ventricle; AA: Atrial appendage; eVariant: Variant underling an eQTL

Together, by expanding the set of associated variants based on high LD (Table 1), overlap with cardiac open chromatin regions (Tables 2 and 3), and prioritizing when known to act as cardiac eQTLs (Table 3), we have generated an unbiased and comprehensive set of candidate causal *cis*-regulatory variants (Supplemental Datasets S1 to S12) underlying the 12 selected QT interval GWAS loci (Figure 1 and Supplemental Figures S1 to S9), which can be evaluated in functional studies.

**Figure 1:**
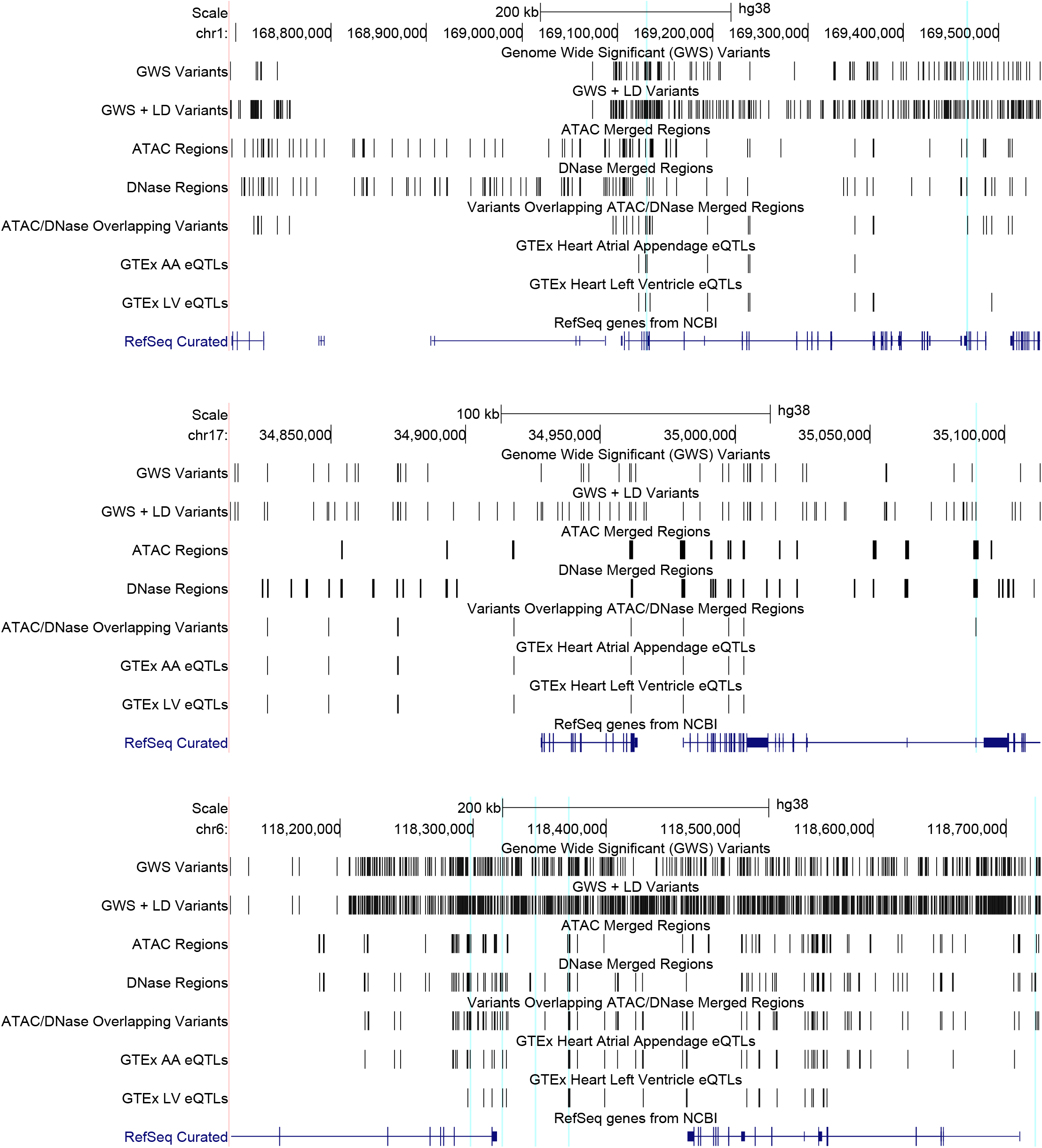
Genomic maps of three representative QT interval associated GWAS loci at *ATP1B1* (*top*), *LIG3* (*middle*) and *PLN* (*bottom*) on chromosomes 1q24.2, 17q12 and 6q22.2-q22.31, respectively, annotated with tracks showing (from top) the genome-wide significant (GWS) variants from the GWAS (GWS Variants); GWS variants plus common variants (minor allele frequency >1%) in high linkage disequilibrium (LD; *r*^2^>0.9) with any of the GWS variants (GWS + LD Variants); merged open chromatin regions from ENCODE ATAC-seq (ATAC Regions) or DNase-seq (DNase Regions) in adult left ventricle tissue samples; associated variants that overlap ATAC-seq and/or DNase-seq left ventricle open chromatin regions (ATAC/DNase Overlapping Variants); open chromatin overlapping variants that are known to be expression quantitative trait loci (eQTLs) in GTEx atrial appendage (AA; GTEx AA eQTLs) or left ventricle (LV; GTEx LV eQTLs) single-tissue *cis*-eQTL data; and the protein-coding genes from NCBI (RefSeq Curated).

## Discussion

Electrocardiographic QT interval, an index of cardiac ventricular de- and re-polarization (35), is a heritable quantitative trait (36). Although being used here as a model complex trait to understand the molecular mechanisms underlying GWAS signals, QT interval is clinically relevant as its prolongation or shortening is associated with an increased risk for cardiovascular morbidity and mortality, largely in the form of cardiac arrhythmias, which can be fatal and lead to sudden cardiac death (37, 38).

Starting with the first report in 2006 (32), GWAS of QT interval have identified dozens of associated loci (20). Like vast majority of typical GWAS (39), lack of genetic diversity is evident in QT interval GWAS as well, which despite efforts to diversify have remained focused on European ancestry subjects (20). One way to address this limitation is to identify the molecular mechanisms underlying existing loci, a long-term goal of this study, which are expected to be shared across ancestries (40). Furthermore, in the present study, we selected 12 QT interval GWAS loci, which have been well replicated across multiple studies (19, 20) and have remained in the top scoring loci based on variance explained (19-22). These loci also harbor genes, which through prior genetic (Mendelian) and biochemical studies are known to directly or indirectly influence cardiomyocyte action potential duration or excitation-coupling, thereby representing critical rate-limiting steps in regulation of ventricular de- and re-polarization.

Our rationale for using nucleosome-free open chromatin for filtering potential *cis*-regulatory causal variants is based on the observation that open chromatin sites are universal hallmarks of all types of gene regulatory elements, including promoters, enhancers, silencers, insulators, and locus control regions (9, 17). With the goal to filter variants in an unbiased and comprehensive manner, we do not utilize additional epigenomic annotations such as specific histone modifications that are correlated with subtypes of regulatory elements (10). Although we have previously generated DNase-seq based open chromatin maps in human adult left ventricle tissue (41), increase in the number of individual samples (biological replicates) available through ENCODE for DNase-seq was the primary driving factor for generating these maps again with increased sensitivity to detect all open chromatin regions. Also, as DNase-seq and ATAC-seq capture open chromatin regions with different efficiencies, leading to ∼35% nonoverlapping distinct peaks (16), generation of ATAC-seq based cardiac open chromatin maps was warranted. Lastly, to maintain high confidence in DNase-seq and ATAC-seq defined open chromatin regions, we only retain those that are observed in two or more biological replicates.

Though, the critical role of genetic variation in gene expression regulation was well established in the pre-GWAS era (12), application of GWAS to complex traits has paralleled the efforts for large-scale eQTL mapping (14). And following the *cis*-regulatory mechanistic hypothesis underlying GWAS, interpretation of GWAS results with eQTL data has been widely used to identify regulatory variants and their target genes at GWAS loci. However, this integration has several challenges, and often leads to false positive discovery of causal genes and functional hypothesis at GWAS loci (42), with a recent report even implying that GWAS mapping and eQTL mapping are systematically biased towards different types of variants (43). Acknowledging these challenges, in our approach we have used overlap with open chromatin regions as the primary filter to identify candidate causal variants, and used colocalization of these filtered variants with known eQTLs as a secondary prioritization step.

Both at the level of LD based expansion of associated candidate variants and open chromatin overlap based variant filtering, we observed limited variability across the 12 GWAS loci, suggesting an underlying uniformity in our approach. Inclusion of high LD common variants led to similar fold-change relative to the number of variants prior to LD-based expansion across the 12 loci (mean: 2.89-fold; median 2.53-fold; range: 1.97-5.15-fold; Table 1), with smallest and largest gains at *LIG3* and *LITAF* loci, respectively. Similarly, on average 15.9% of candidate variants overlapped cardiac open chromatin regions across the 12 loci (median: 14.1%; range: 5.0%-31.6%; Table 3), with *KCNJ2* (5.0%) and *RNF207* (31.6%) being the outliers. Overall, using a simple yet impactful bioinformatic approach, we have generated a comprehensive list of candidate causal *cis*-regulatory variants at the 12 selected QT interval GWAS loci, all set to be evaluated in regulatory functional experiments.

## Acknowledgements

This work was supported by funds from National Institutes of Health (NIH), USA grant R01 HL158901 (to A.K.) and McGovern Medical School UTHealth (to A.K.). The authors acknowledge the Texas Advanced Computing Center at the University of Texas at Austin for providing the computing and storage resources. We are thankful to Dongwon Lee (Boston Children’s Hospital) for assistance with processing of BAM files and peak calling.

## Author Contributions

A.K. conceived and designed the study. S.K. and A.K. conducted the analyses and wrote the manuscript.

