## Supplemental Tables and Figures for "Identification of candidate causal *cis*-regulatory variants underlying electrocardiographic QT interval GWAS loci"

**This PDF file includes:**

Table S1
Figures S1 to S9

**Other supplementary materials for this manuscript include the following:**

Datasets S1 to S14

**Table S1:** ENCODE human adult left ventricle chromatin accessibility experiments

| Assay | Experiment ID | BAM File ID | Used | Notes from ENCODE Data Coordination Center audits |
|---|---|---|---|---|
| DNase-seq | ENCSR299QGI | ENCFF631HMZ | Yes | N/A |
| DNase-seq | ENCSR395HAE | ENCFF154ILG | Yes | N/A |
| DNase-seq | ENCSR747SEU | ENCFF041ZOJ | Yes | N/A |
| DNase-seq | ENCSR222CLC | ENCFF284XQW | Yes | N/A |
| DNase-seq | ENCSR805EGZ | ENCFF221NLC | Yes | N/A |
| DNase-seq | ENCSR598RVJ | ENCFF682TBE | Yes | N/A |
| DNase-seq | ENCSR735NTV | ENCFF558NDU | Yes | N/A |
| ATAC-seq | ENCSR286STX | ENCFF440WVI | Yes | N/A |
| ATAC-seq | ENCSR899YEP | ENCFF771XJN | Yes | N/A |
| ATAC-seq | ENCSR310RJN | ENCFF804KIW | Yes | N/A |
| ATAC-seq | ENCSR846VPV | ENCFF456PVX | Yes | N/A |
| ATAC-seq | ENCSR925LGW | ENCFF827WPC | Yes | N/A |
| ATAC-seq | ENCSR399OSE | ENCFF160LDI | Yes | N/A |
| DNase-seq | ENCSR070CMW | ENCFF533LLW | No | Noncompliant SPOT score |
| ATAC-seq | ENCSR204PZT | ENCFF919KRC | No | Noncompliant FRiP score |
| ATAC-seq | ENCSR848TMJ | ENCFF704VYQ | No | Noncompliant NRF score |
| ATAC-seq | ENCSR451JSB | ENCFF987YOH | No | Noncompliant FRiP score |
| ATAC-seq | ENCSR225FZH | ENCFF670QFL | No | Noncompliant FRiP score |
| ATAC-seq | ENCSR769DGC | ENCFF627RHW | No | Noncompliant FRiP score |
| ATAC-seq | ENCSR745CGG | ENCFF420QMN | No | Noncompliant FRiP score |
| ATAC-seq | ENCSR593YFB | ENCFF283TUN | No | Noncompliant read depth |
| ATAC-seq | ENCSR117PYB | ENCFF713UOC | No | Noncompliant FRiP score |
| ATAC-seq | ENCSR851EBF | ENCFF097EXD | No | Noncompliant FRiP score and read depth |

N/A: Not applicable.

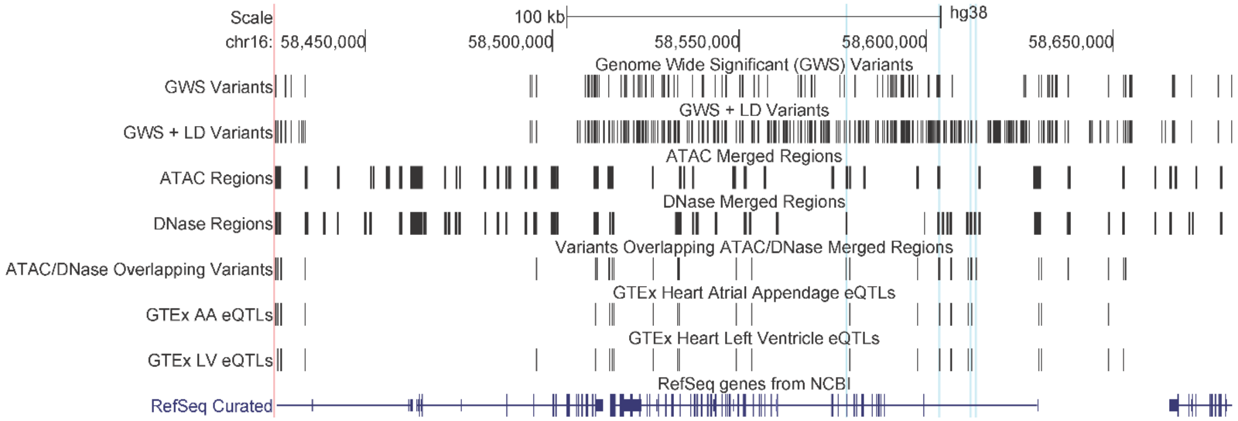


**Figure S1:** Genomic map of QT interval associated *CNOT1* GWAS locus on chromosome 16q21, annotated with tracks showing (from top) the genome wide significant (GWS) variants from the GWAS (GWS Variants); GWS variants plus common variants (minor allele frequency >1%) in high linkage disequilibrium (LD; $r^2>0.9$) with any of the GWS variants (GWS + LD Variants); merged open chromatin regions from ENCODE ATAC-seq in adult left ventricle tissue samples (ATAC Regions); merged open chromatin regions from ENCODE DNase-seq in adult left ventricle tissue samples (DNase Regions); associated variants that overlap ATAC-seq and/or DNase-seq left ventricle open chromatin regions (ATAC/DNase Overlapping Variants); open chromatin overlapping variants that are known to be expression quantitative trait loci (eQTLs) in GTEx atrial appendage (AA) single-tissue *cis*-eQTL data (GTEx AA eQTLs); open chromatin overlapping variants that are known to be eQTLs in GTEx left ventricle (LV) single-tissue *cis*-eQTL data (GTEx LV eQTLs); and the protein-coding genes from NCBI (RefSeq Curated).

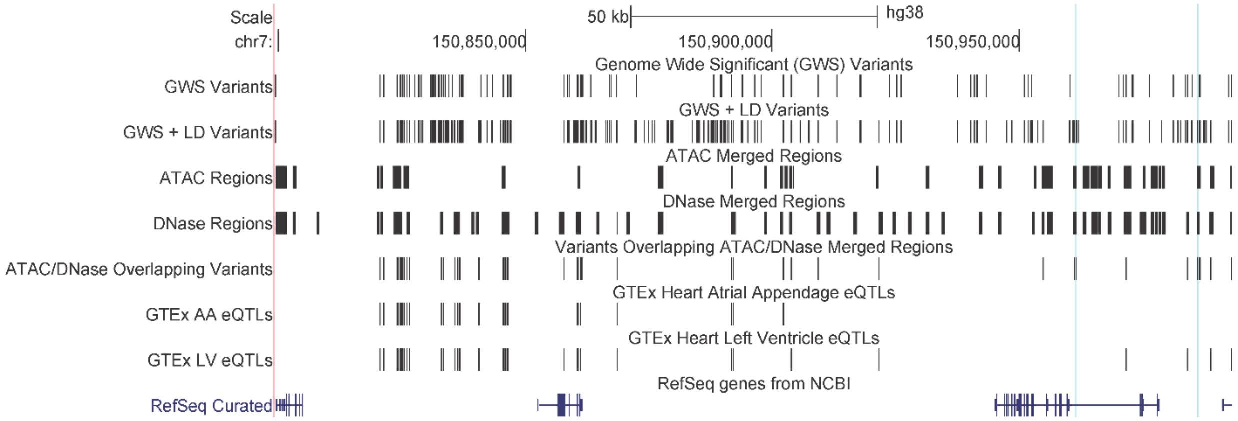


**Figure S2:** Genomic map of QT interval associated *KCNH2* GWAS locus on chromosome 7q36.1, annotated with tracks showing (from top) the genome wide significant (GWS) variants from the GWAS (GWS Variants); GWS variants plus common variants (minor allele frequency >1%) in high linkage disequilibrium (LD; $r^2>0.9$) with any of the GWS variants (GWS + LD Variants); merged open chromatin regions from ENCODE ATAC-seq in adult left ventricle tissue samples (ATAC Regions); merged open chromatin regions from ENCODE DNase-seq in adult left ventricle tissue samples (DNase Regions); associated variants that overlap ATAC-seq and/or DNase-seq left ventricle open chromatin regions (ATAC/DNase Overlapping Variants); open chromatin overlapping variants that are known to be expression quantitative trait loci (eQTLs) in GTEx atrial appendage (AA) single-tissue *cis*-eQTL data (GTEx AA eQTLs); open chromatin overlapping variants that are known to be eQTLs in GTEx left ventricle (LV) single-tissue *cis*-eQTL data (GTEx LV eQTLs); and the protein-coding genes from NCBI (RefSeq Curated).

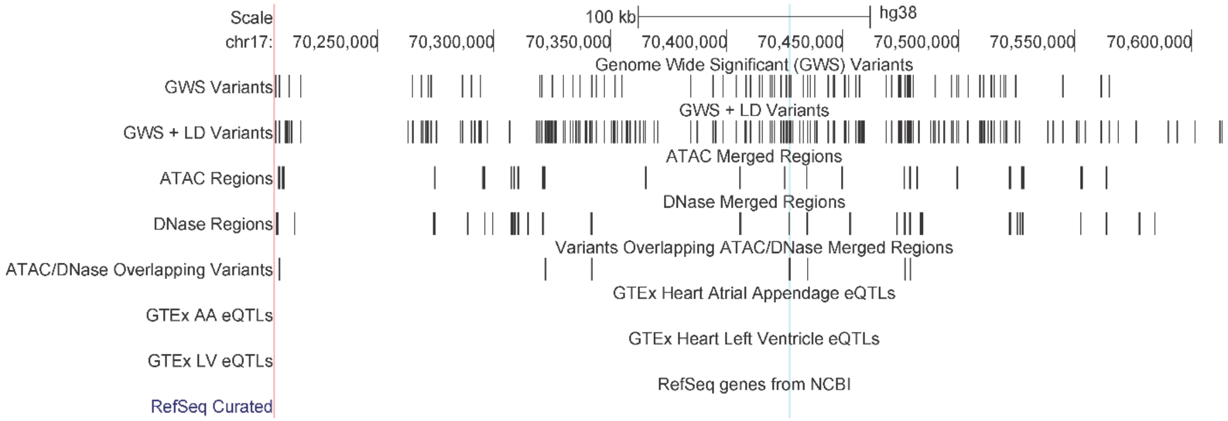


**Figure S3:** Genomic map of QT interval associated *KCNJ2* GWAS locus on chromosome 17q24.3, annotated with tracks showing (from top) the genome wide significant (GWS) variants from the GWAS (GWS Variants); GWS variants plus common variants (minor allele frequency >1%) in high linkage disequilibrium (LD; $r^2$>0.9) with any of the GWS variants (GWS + LD Variants); merged open chromatin regions from ENCODE ATAC-seq in adult left ventricle tissue samples (ATAC Regions); merged open chromatin regions from ENCODE DNase-seq in adult left ventricle tissue samples (DNase Regions); associated variants that overlap ATAC-seq and/or DNase-seq left ventricle open chromatin regions (ATAC/DNase Overlapping Variants); open chromatin overlapping variants that are known to be expression quantitative trait loci (eQTLs) in GTEx atrial appendage (AA) single-tissue *cis*-eQTL data (GTEx AA eQTLs); open chromatin overlapping variants that are known to be eQTLs in GTEx left ventricle (LV) single-tissue *cis*-eQTL data (GTEx LV eQTLs); and the protein-coding genes from NCBI (RefSeq Curated).

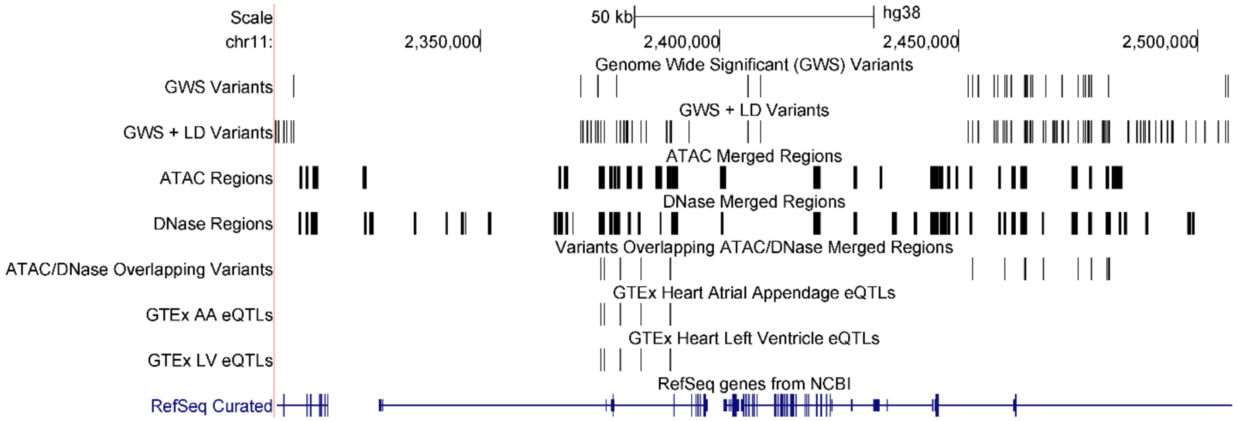


**Figure S4:** Genomic map of QT interval associated *KCNQ1* GWAS locus on chromosome 11p15.5, annotated with tracks showing (from top) the genome wide significant (GWS) variants from the GWAS (GWS Variants); GWS variants plus common variants (minor allele frequency >1%) in high linkage disequilibrium (LD; $r^2$>0.9) with any of the GWS variants (GWS + LD Variants); merged open chromatin regions from ENCODE ATAC-seq in adult left ventricle tissue samples (ATAC Regions); merged open chromatin regions from ENCODE DNase-seq in adult left ventricle tissue samples (DNase Regions); associated variants that overlap ATAC-seq and/or DNase-seq left ventricle open chromatin regions (ATAC/DNase Overlapping Variants); open chromatin overlapping variants that are known to be expression quantitative trait loci (eQTLs) in GTEx atrial appendage (AA) single-tissue *cis*-eQTL data (GTEx AA eQTLs); open chromatin overlapping variants that are known to be eQTLs in GTEx left ventricle (LV) single-tissue *cis*-eQTL data (GTEx LV eQTLs); and the protein-coding genes from NCBI (RefSeq Curated).

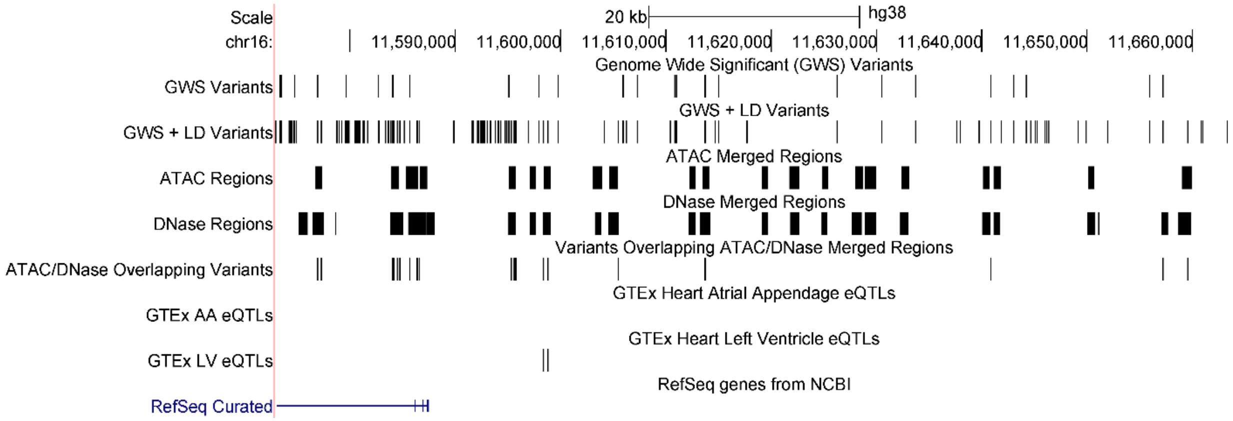


**Figure S5:** Genomic map of QT interval associated *LITAF* GWAS locus on chromosome 16p13.13, annotated with tracks showing (from top) the genome wide significant (GWS) variants from the GWAS (GWS Variants); GWS variants plus common variants (minor allele frequency >1%) in high linkage disequilibrium (LD; $r^2$>0.9) with any of the GWS variants (GWS + LD Variants); merged open chromatin regions from ENCODE ATAC-seq in adult left ventricle tissue samples (ATAC Regions); merged open chromatin regions from ENCODE DNase-seq in adult left ventricle tissue samples (DNase Regions); associated variants that overlap ATAC-seq and/or DNase-seq left ventricle open chromatin regions (ATAC/DNase Overlapping Variants); open chromatin overlapping variants that are known to be expression quantitative trait loci (eQTLs) in GTEx atrial appendage (AA) single-tissue *cis*-eQTL data (GTEx AA eQTLs); open chromatin overlapping variants that are known to be eQTLs in GTEx left ventricle (LV) single-tissue *cis*-eQTL data (GTEx LV eQTLs); and the protein-coding genes from NCBI (RefSeq Curated).

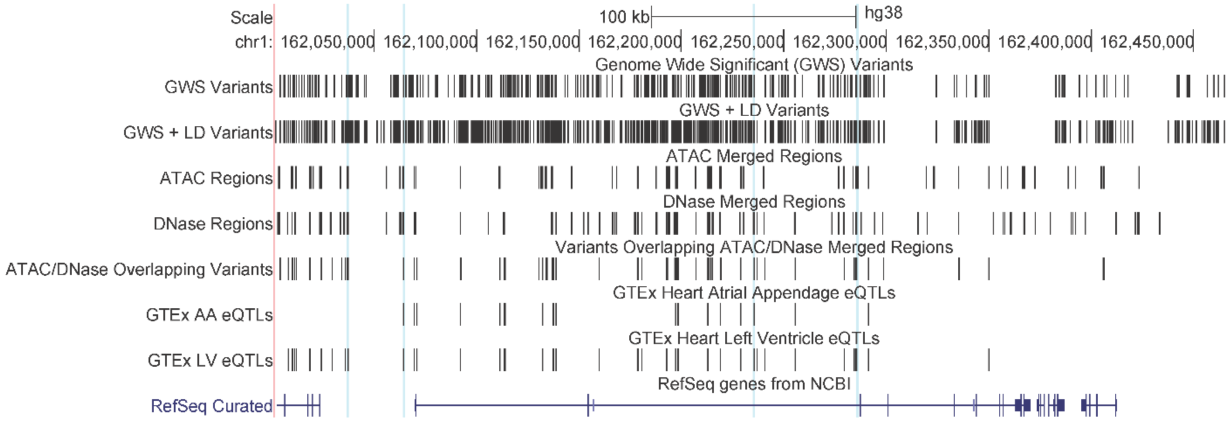


**Figure S6:** Genomic map of QT interval associated *NOS1AP* GWAS locus on chromosome 1q23.3, annotated with tracks showing (from top) the genome wide significant (GWS) variants from the GWAS (GWS Variants); GWS variants plus common variants (minor allele frequency >1%) in high linkage disequilibrium (LD; $r^2>0.9$) with any of the GWS variants (GWS + LD Variants); merged open chromatin regions from ENCODE ATAC-seq in adult left ventricle tissue samples (ATAC Regions); merged open chromatin regions from ENCODE DNase-seq in adult left ventricle tissue samples (DNase Regions); associated variants that overlap ATAC-seq and/or DNase-seq left ventricle open chromatin regions (ATAC/DNase Overlapping Variants); open chromatin overlapping variants that are known to be expression quantitative trait loci (eQTLs) in GTEx atrial appendage (AA) single-tissue *cis*-eQTL data (GTEx AA eQTLs); open chromatin overlapping variants that are known to be eQTLs in GTEx left ventricle (LV) single-tissue *cis*-eQTL data (GTEx LV eQTLs); and the protein-coding genes from NCBI (RefSeq Curated).

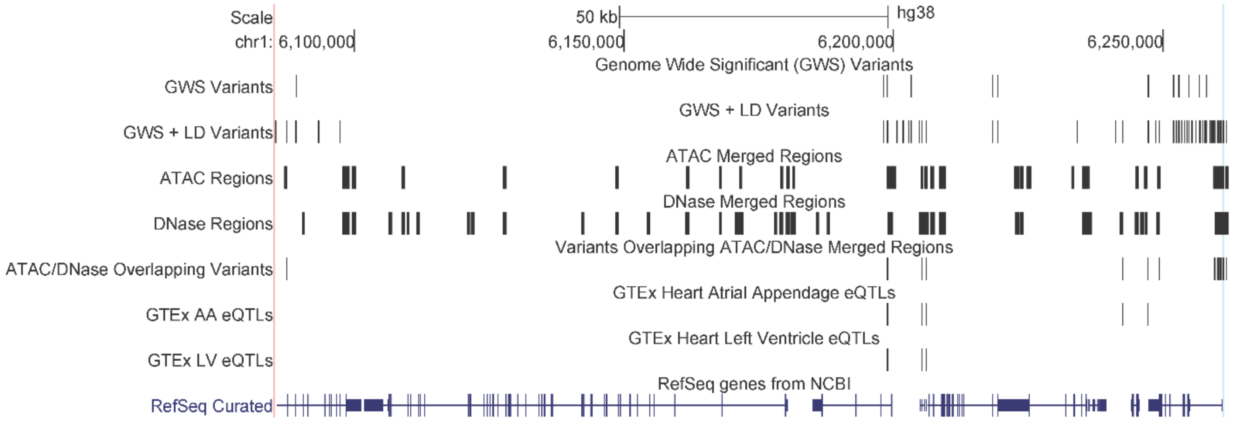


**Figure S7:** Genomic map of QT interval associated *RNF207* GWAS locus on chromosome 1p36.31, annotated with tracks showing (from top) the genome wide significant (GWS) variants from the GWAS (GWS Variants); GWS variants plus common variants (minor allele frequency >1%) in high linkage disequilibrium (LD; $r^2$>0.9) with any of the GWS variants (GWS + LD Variants); merged open chromatin regions from ENCODE ATAC-seq in adult left ventricle tissue samples (ATAC Regions); merged open chromatin regions from ENCODE DNase-seq in adult left ventricle tissue samples (DNase Regions); associated variants that overlap ATAC-seq and/or DNase-seq left ventricle open chromatin regions (ATAC/DNase Overlapping Variants); open chromatin overlapping variants that are known to be expression quantitative trait loci (eQTLs) in GTEx atrial appendage (AA) single-tissue *cis*-eQTL data (GTEx AA eQTLs); open chromatin overlapping variants that are known to be eQTLs in GTEx left ventricle (LV) single-tissue *cis*-eQTL data (GTEx LV eQTLs); and the protein-coding genes from NCBI (RefSeq Curated).

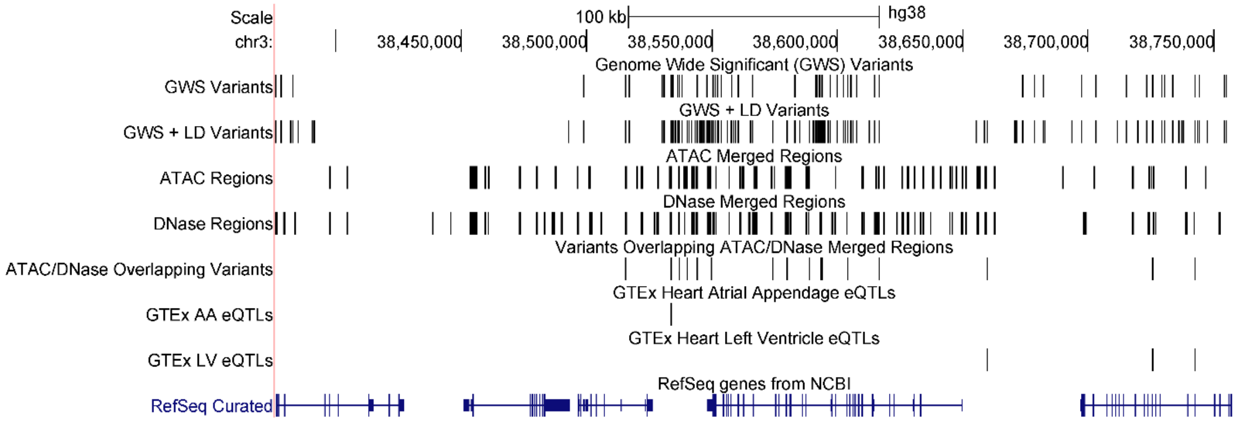


**Figure S8:** Genomic map of QT interval associated *SCN5A* GWAS locus on chromosome 3p22.2, annotated with tracks showing (from top) the genome wide significant (GWS) variants from the GWAS (GWS Variants); GWS variants plus common variants (minor allele frequency >1%) in high linkage disequilibrium (LD; $r^2$>0.9) with any of the GWS variants (GWS + LD Variants); merged open chromatin regions from ENCODE ATAC-seq in adult left ventricle tissue samples (ATAC Regions); merged open chromatin regions from ENCODE DNase-seq in adult left ventricle tissue samples (DNase Regions); associated variants that overlap ATAC-seq and/or DNase-seq left ventricle open chromatin regions (ATAC/DNase Overlapping Variants); open chromatin overlapping variants that are known to be expression quantitative trait loci (eQTLs) in GTEx atrial appendage (AA) single-tissue *cis*-eQTL data (GTEx AA eQTLs); open chromatin overlapping variants that are known to be eQTLs in GTEx left ventricle (LV) single-tissue *cis*-eQTL data (GTEx LV eQTLs); and the protein-coding genes from NCBI (RefSeq Curated).

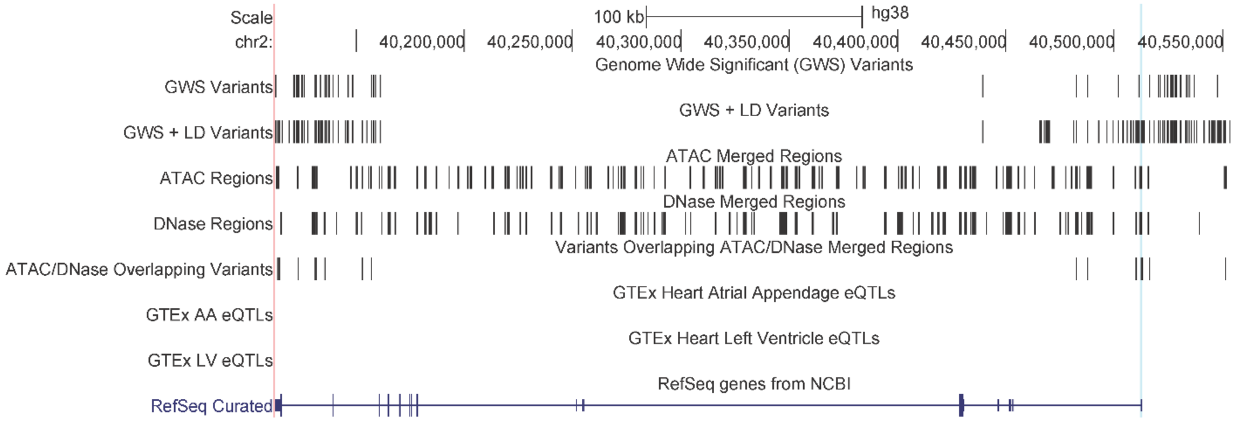


**Figure S9:** Genomic map of QT interval associated *SLC8A1* GWAS locus on chromosome 2p22.1, annotated with tracks showing (from top) the genome wide significant (GWS) variants from the GWAS (GWS Variants); GWS variants plus common variants (minor allele frequency >1%) in high linkage disequilibrium (LD; $r^2$>0.9) with any of the GWS variants (GWS + LD Variants); merged open chromatin regions from ENCODE ATAC-seq in adult left ventricle tissue samples (ATAC Regions); merged open chromatin regions from ENCODE DNase-seq in adult left ventricle tissue samples (DNase Regions); associated variants that overlap ATAC-seq and/or DNase-seq left ventricle open chromatin regions (ATAC/DNase Overlapping Variants); open chromatin overlapping variants that are known to be expression quantitative trait loci (eQTLs) in GTEx atrial appendage (AA) single-tissue *cis*-eQTL data (GTEx AA eQTLs); open chromatin overlapping variants that are known to be eQTLs in GTEx left ventricle (LV) single-tissue *cis*-eQTL data (GTEx LV eQTLs); and the protein-coding genes from NCBI (RefSeq Curated).
